# sHSP-induced oligomerization enhances mammalian transcription factor potency

**DOI:** 10.64898/2026.11.12.766240

**Authors:** Tyler J. Buckley, Mannix J. Burns, Hanna Liliom, Su May Lei Soe, Brandon Boone, Shuya Wang, Yan He, Boey Li, Maya Hanley, Sophia Sauceda, Suhua Feng, Débora R. Sobreira, Steven E. Jacobsen

## Abstract

Recently, a non-canonical role for small heat shock proteins (sHSPs) was identified, wherein sHSP-induced oligomerization drives targeted genomic accumulation of an endogenous silencing complex. Here we repurpose this property to develop sHSPs as compact, tunable tags for transcriptional engineering. Screening sHSPs across the tree of life, we show that sHSPs fused to transcription factors (TFs) can increase on-target genomic occupancy, recruit sHSP-tagged proteins, and amplify target-gene activation in mammalian cells. Distinct sHSPs vary in levels of transcriptional occupancy and output, providing a means to tune TFs activity while maintaining targeting specificity. This work establishes sHSPs as a new class of compact (∼20kDa), tunable, activity-enhancing tags with broad potential for transcriptional engineering and synthetic biology.

## Introduction

A central goal in transcriptional engineering is to tune regulatory activity at specific genomic loci and the recruitment of functional proteins while maintaining targeting specificity. Transcription factors (TFs) and CRISPR based systems can be used to target specific genomic sites, but the magnitude of their effects is often limited by genomic occupancy and how much additional functional proteins can be recruited there. Existing systems to increase local protein concentration, including SunTag and coiled-coil-based recruitment, often relies on large or multi-component interactions^1,2^. Therefore, a compact system to enhance and tune regulatory activity while preserving native targeting specificity of fused proteins could provide a useful tool for transcriptional engineering.

Alpha-crystalline domain (ACD) containing small heat shock proteins (sHSPs) are compact, evolutionarily conserved chaperones that keep partner proteins stably folded during periods of cellular stress^3,4^. Under homeostatic conditions, most sHSPs are sequestered into large inert oligomers^5^ mediated between their conserved ACD domain and C-terminal IXI-V-like motif.

Importantly, oligomerization varies among sHSPs, ranging from dimers to large oligomeric complexes. This natural diversity, with their small size, suggests that sHSPs could be used in targeted protein accumulation.

Recently, a non-canonical role for sHSPs was identified in *Arabidopsis thaliana*, wherein sHSP-induced oligomerization drives targeted genomic accumulation of an endogenous DNA methylation-dependent silencing complex called the MBD5/6 complex^6,7,8^. In this system, methyl-DNA binding proteins MBD5 and MBD6 recruit two sHSPs, ACD15 and ACD21, which are required for accumulation of complex members and can form large biomolecular condensate that are visible as nuclear foci. Foci formation is abolished in *acd15 acd21* mutants, confirming these sHSPs are required for complex accumulation^6^.

We hypothesized that sHSP oligomerization could be repurposed by directly fusing sHSPs to TFs to locally concentrate engineered regulatory proteins at defined genomic sites without altering their intrinsic targeting specificity (**Fig. 1A**). Furthermore, interactions between identical sHSPs could allow additional sHSP-tagged proteins to be recruited to TF-bound loci.

**Figure 1:**
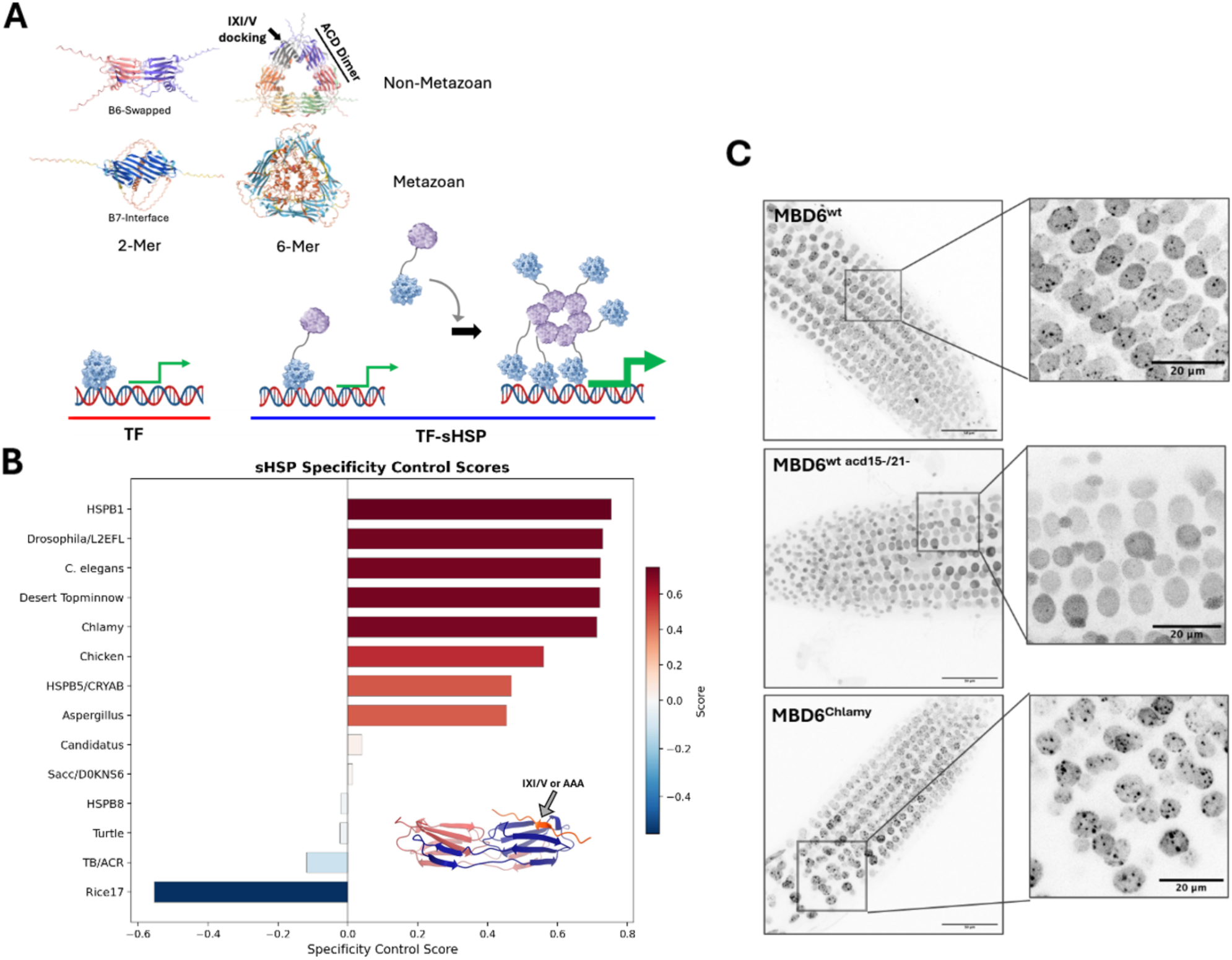
Oligomerization of exogenous sHSPs is sufficient to recapitulate genomic accumulation of an endogenous silencing complex. **A)** Schematic of sHSP dimer and hexamer organization (TOP) and sHSP-TF fusion system (Bottom). **B)** AlphaFold3 predicted docking score between the canonical IXI/V motif and the ACD domain of each synthetically screened sHSP **C)** Representative images of immunofluorescence microscopy of MBD6-YFP in WT and *acd15/21* plants, and MBD6-Chlamy-YFP in *acd15/21* plants. Color is scale is set to black/white and inverted.

Here, we screened sHSPs spanning the tree of life and tested their ability as transcriptional engineering tags. We show that fusion of an sHSP to Myc increases its accumulation at endogenous Myc-binding sites and amplifies Myc-associated transcriptional responses in mammalian cells We further show that sHSP-tagged proteins can be recruited to TF-bound genomic loci through interactions with a TF-sHSP fusion. Using a panel of sHSPs spanning a range of oligomerization capacities, we show that distinct sHSPs have different levels of transcriptional activity, providing a means to tune TF potency. Finally, fusion of an sHSP to the pioneer TF NGN2 enhanced transcriptional activation in primary human fibroblasts. Together, these results establish sHSPs as compact, tunable, activity-enhancing tags for transcriptional engineering.

## Results

To prioritize candidate sHSPs, we performed an AlphaFold 3 based docking assay on a diverse range of sHSPs spanning the tree of life. This analysis tested the docking specificity of the predicted multimerization interaction between the C-terminal IXI/V-like motif and the Alpha Crystallin Domain of each sHSP, together with an AAA motif substitution control (**Fig. 1B, Methods**)^9^. In line with previous characterizations, HSPB1 and HSPB5, which are known to form large oligomers, had high specificity scores, whereas dimer-only forming HSPB8 had a low specificity score^10^. From this analysis we chose a sHSP from *Chlamydomonas reinhardtii,* CHLRE_07g318800v5, (Chlamy) with a high specificity score, and that was therefore predicted to highly multimerize. We then tested whether Chlamy could restore MBD5/6 complex accumulation and foci formation in *acd15 acd21* plants by fusing it to MBD6 in transgenic Arabidopsis plants and performed immunofluorescence microscopy. We observed that fusion with Chlamy was sufficient to restore nuclear foci formation of the MBD5/6 complex (**Fig. 1C**), indicative of genomic on-target oligomerization.

Next, to test whether sHSP oligomerization could increase transcription factor (TF) occupancy in mammalian chromatin, we fused Chlamy to the C terminus of full-length Myc. Using doxycycline-inducible PiggyBac constructs in HEK293T cells, we compared IgG control, Myc, and Myc-Chlamy binding profiles 24 hours after induction by CUT&RUN. Myc-Chlamy retained the genomic specificity of Myc but exhibited amplified and broadened signal (**Fig. 2A**), even though transcript abundance of Myc and Myc-Chlamy were similar (**Suppl. Fig. 1**). Across the 52,089 merged MACS2-called peaks across all samples, Myc and Myc-Chlamy were enriched at the same genomic loci (Pearson correlation = 0.70, p < 2.2e-16). The spike-in normalized signal at peak centers (+/-250bp) was higher for Myc-Chlamy than Myc at 87% of peaks (median of 1.79-fold-change, p = 0.0002) and the average binding profile was broader (Myc FWHM mean 367+/-29bp, Myc-Chlamy FWHM mean 433+/-29bp, p = 0.047). Importantly, peaks called in only one treatment group showed signal enrichment in the other group, indicating that fusion with Chlamy did not redirect Myc to unrelated binding sites, but increased its ability to accumulate at endogenous targets.

**Figure 2:**
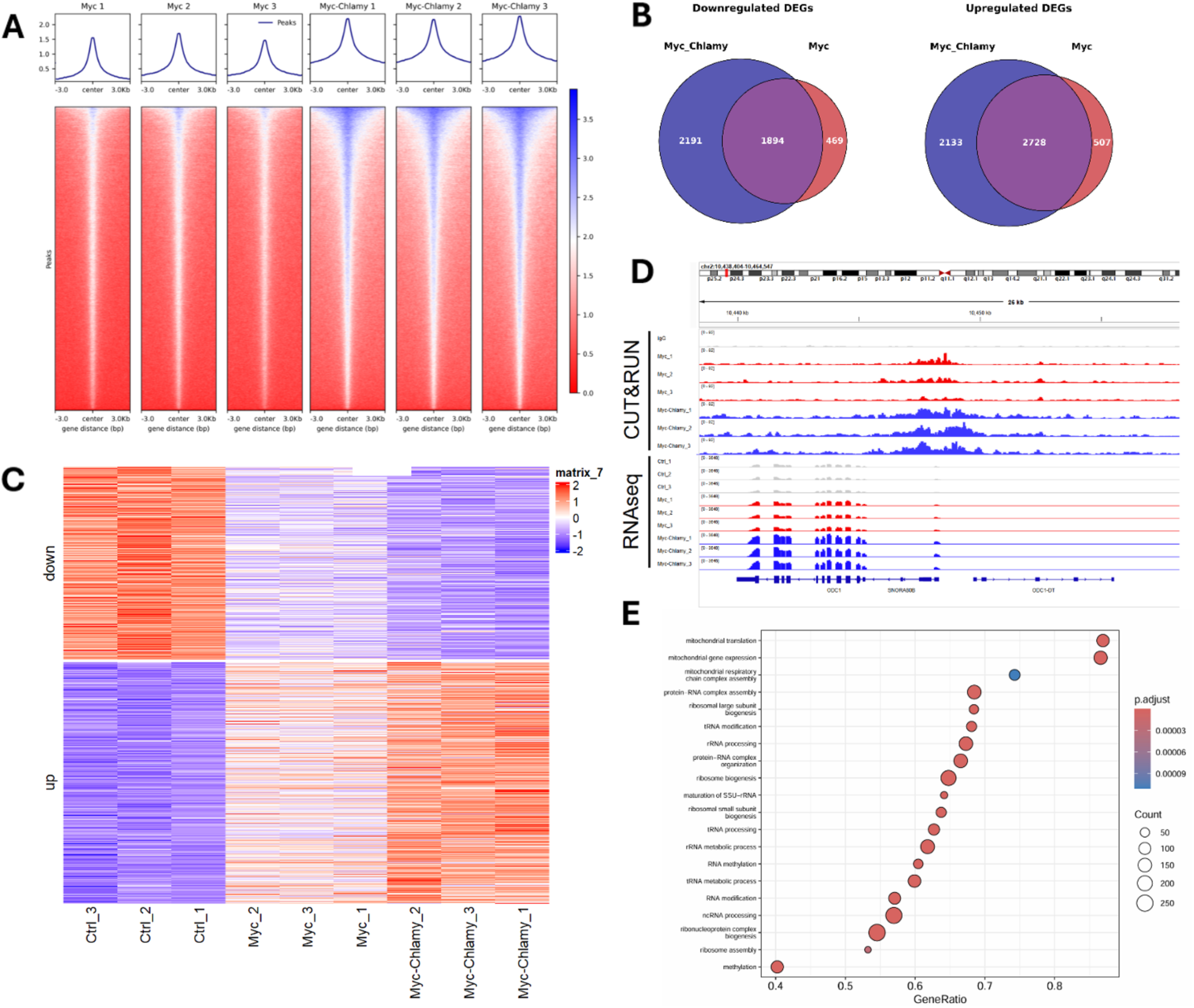
sHSP-fusion enhances the genomic association and potency of a Transcription Factor. **A)** IgG-normalized CUT&RUN heatmap showing binding signal for Myc and Myc-Chlamy at peaks identified by macs2 callpeak in either group **B)** Venn diagram displaying differentially expressed down-(left) or up-regulated (right) genes in Myc or Myc-Chlamy compared to control (DESeq2, p.adj < 0.05) **C)** RNAseq heatmap of differentially expressed genes (DESeq2, p.adj < 0.05) in Myc and Myc-Chlamy overexpressing cells compared to control **D)** Integrated Genome Browser track visualization of spike-in normalized CUT&RUN peaks (top) and RNAseq signal (bottom) at the ODC1 gene, a canonical Myc target **E)** Biological Process Gene Set Enrichment Analysis for differentially expressed genes in Myc and Myc-Chlamy compared to control (DESeq2, p.adj < 0.05).

To determine whether this sHSP-driven genomic accumulation results in increased TF activity, we performed RNA-seq on the same set of inducible GFP control, Myc, and Myc-Chlamy HEK293T cells as previously described. Similar to our CUT&RUN results, we observed strongly overlapping signatures of differentially expressed genes in Myc and Myc-Chlamy compared to GFP control (**Fig. 2B**). We found that genes impacted by Myc-Chlamy trended towards a greater magnitude of change than Myc, with more differentially expressed genes detected in Myc-Chlamy (4,861 up, 4,085 down, p.adj < 0.05) compared to Myc (3,235 up, 2,363 down, p.adj < 0.05). When visualizing differentially expressed genes (DEGs) between both Myc and Myc-Chlamy together compared to control, we observed a clear stepwise change in expression that was found across replicates from the GFP control, to Myc, to Myc-Chlamy (**Fig. 2C**), for example as shown at the representative known Myc target ODC1^11^ (**Fig. 2D**). In conjunction with the binding data, these data suggest that Chlamy fusion induces on-target oligomerization to enhance the potency of Myc. To investigate the processes impacted by Myc and Myc-Chlamy, we performed Gene Set Enrichment Analysis (GSEA) and found enrichment for pathways related to translation and metabolism, consistent with previous characterizations of Myc function (**Fig. 2E**)^12^. We investigated the genes detected as differentially expressed in Myc-Chlamy but not Myc and found that they also trended upwards in Myc compared to control, consistent with the sHSP conferring a larger magnitude of effect (**Suppl. Fig. 2**). Overall, this transcriptomic data shows that the fusion of an sHSP to a TF enhances the potency of that TF.

To validate that this increased sHSP-driven activity can be recapitulated in other cell types and with different TF fusions, we fused Chlamy with the pioneering TF NGN2, and performed RNA-seq of GFP control, NGN2, and NGN2-Chlamy in primary adult human dermal fibroblasts. Consistent with an activity-enhancing effect, we observed that NGN2-Chlamy upregulated shared target genes more than NGN2 alone (**Suppl. Fig. 3**). These results utilizing a pioneer factor in primary human cells provides a relevant basis for future work employing sHSP-fusions for cellular reprogramming applications.

The TF-Chlamy fusions are predicted to form target localized multimers capable of recruiting additional effector protein-Chlamy fusions to transcription factor-bound loci. To test this, we generated a dual-tag construct co-expressing Myc-Chlamy-FLAG and Chlamy-HA, reasoning that Chlamy-HA, which lacks a DNA-binding domain, should accumulate on chromatin only through oligomerization with genome-bound Myc-Chlamy-FLAG (**Fig. 3A**).

**Figure 3:**
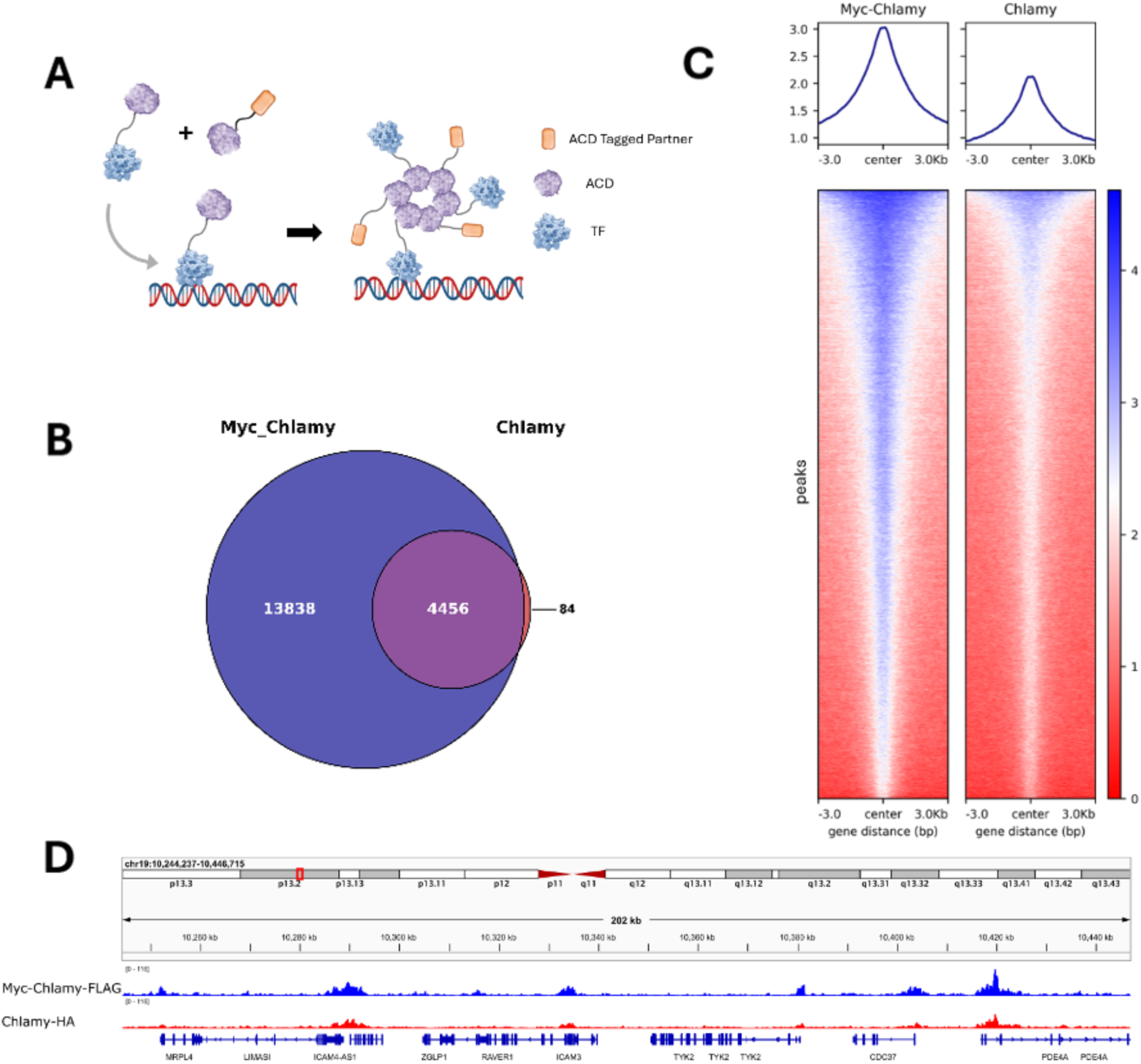
sHSPs can be multiplexed for protein recruitment. **A)** Schematic of the dual-tag sHSP recruitment system **B)** Venn diagram showing overlap between macs2 callpeak peaks called for Myc-Chlamy-FLAG and Chlamy-HA **C)** IgG-normalized CUT&RUN heatmap showing binding signal for Myc and Myc-Chlamy at peaks identified as significant in either group **D)** Integrated Genome Browser track visualization of spike-in normalized CUT&RUN peaks for Myc-Chlamy-FLAG and Chlamy-HA across a range of binding sites.

Indeed, CUT&RUN showed that the co-expressed Chlamy-HA was present at Myc-Chlamy peaks (**Fig. 3B,3C,3D**). These results support a model in which ACD-containing sHSP fusions form target-localized oligomeric assemblies which should allow for the possibility of recruiting additional tagged regulatory proteins to endogenous transcription factor binding sites.

To expand the sHSP toolkit and gain better insight into the impact of oligomerization on TF potency, we harnessed a wide catalog of sHSPs that are naturally occurring across the tree of life, from archaea to mammals, and are known or predicted to have varying oligomerization capacities^1^. We selected a set of sHSPs spanning the predictive oligomerization specificity score range and tested different Myc-sHSP fusions to determine the impact different sHSPs on TF-activity enhancement in HEK293T cells, comparing GFP control, Myc, or Myc-sHSP fusions **(Fig. 4A, Supp. Table 1**). We observed a gradient effect of Myc-sHSP fusions across Myc target genes (**Fig. 4A, 4B**). As expected, there was a high correlation (Pearson correlation = 0.725, p = 0.0033) between sHSP predicted specificity score and sHSP-fusion potency, which was defined as the median upregulated gene z-score value (**Fig 4C**). This indicates that there is a positive relationship between an sHSPs oligomerization capacity and the TF-potency effect that it induces. These results provide evidence that different sHSPs can thus be used to tune TF-activity.

**Figure 4:**
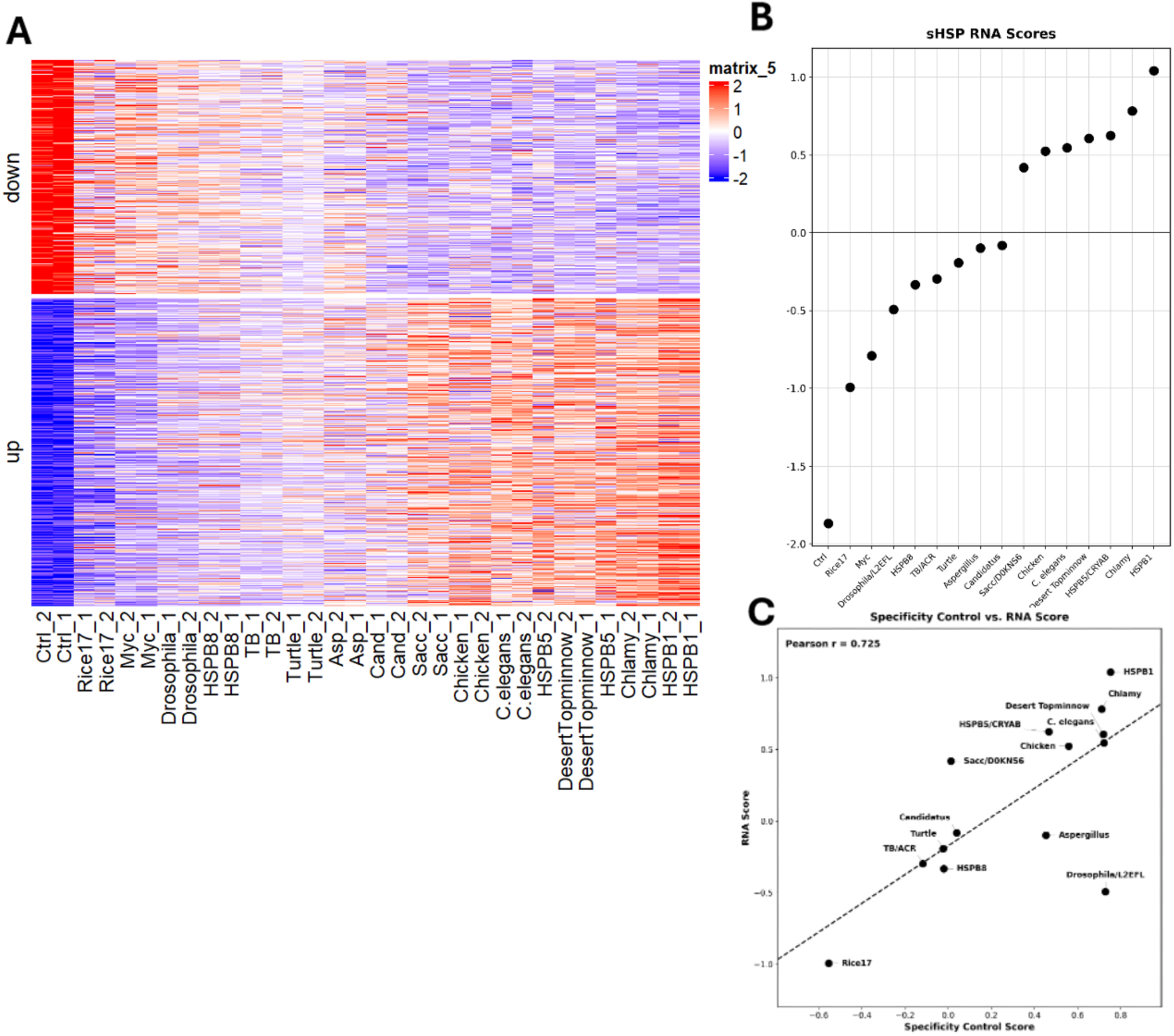
Generation of an sHSP library with varying conferred potency. **A)** RNA heatmap of differentially expressed (DESeq2, p.adj < 0.05) in all Myc-transgenic cells compared to control **B)** Plot displaying a proxy metric for the sHSP-conferred transcriptional potency effect, defined as the median Myc-transgenic upregulated DEG z-score derived from the same data as panel A **C)** Correlation scatterplot between predicted docking specificity score (AlphaFold3, Specificity Score = WT_score - AAA_ctrl_score) and RNA Score, as seen in panel B.

## Discussion

Here we show that an oligomerization-driven mechanism first identified in an endogenous plant silencing complex can be repurposed as a genetically encoded tag for enhancing and tuning TF activity in mammalian cells. Fusion of ACD-containing sHSPs to Myc and to the pioneer factor NGN2 increased genomic accumulation and amplified downstream transcriptional output without altering the intrinsic targeting specificity of the fused TF. Distinct sHSPs also demonstrated graded levels of potency that tracked with their predicted oligomerization capacity. Together with sHSP-tagged proteins co-recruiting at TF-bound loci, these results establish sHSPs as compact tag for transcriptional engineering with a substantially smaller size as existing systems.

Mechanistically, our findings are consistent with a model in which sHSP-driven oligomerization increases the local TF concentration on endogenous binding sites, rather than redirecting them to new loci. Myc-Chlamy retained the genomic specificity of Myc, and peaks unique to either fusion protein were detectably bound by the other, arguing against non-specific binding sites. Instead, Myc-Chlamy occupancy was both higher in magnitude and broader in width than Myc alone, a signature of the spreading and increased local density associated with condensate-like assemblies, and consistent with the oligomerization-dependent nuclear foci we have observed for ACD-containing sHSPs in plants.

Across a panel of sHSPs tested, the magnitude of transcriptional upregulation associated with each sHSPs predicted ACD-IXI/V docking specificity, a prediction for its native oligomerization capacity. This relationship indicates that oligomerization itself is tunable. Rather than choosing between a binary ‘on’ or ‘off’ accumulation tag, sHSP identity can be used as a dial of the degree of TF potentiation to the given application. Because the docking specificity score was calculated from AlphaFold3 structural predictions, without any measurement of oligomeric state, this correlation also suggests that additional computationally predicted or designed ACD variants could be screened to further expand the range of tunable outputs.

The ability of sHSP-tagged proteins to be recruited to TF-bound loci through Chlamy-Chlamy interactions suggests that this may be applied with a higher client number. In principle, co-expression of a TF-sHSP fusion with additional sHSP-tagged effectors, such as chromatin modifiers, additional transcriptional activators, or reporters, could allow assembly of multi-component regulatory condensates at defined genomic loci using a single tag. We also show that sHSP fusions enhanced the activity of two distinct TFs, a highly expressed, ubiquitously active TF, Myc, and a pioneer factor, NGN2, that must first engage closed chromatin, in two different cell types. This supports a model where this system is not restricted to a single class of DNA-binding protein or chromatin context and could be adapted towards applications in which the intrinsic activity of a TF, or of a targeted synthetic effector is limited.

In this study, we ranked our sHSP library based on transcriptomic activity rather than genomic occupancy for every family member, and future work should confirm that occupancy and transcriptional outcomes remain coupled across a wider range of sHSPs and target genes. sHSPs also act as native molecular chaperones, and large assemblies could in principle be acting on other endogenous targets or recruiting other clients. While we did not observe any evidence of this, we believe that having a wide range of sHSP tags from diverse origins will limit any off-target chaperone activity in your system of choice.

In summary, this work repurposes the non-canonical role of sHSP oligomerization into a compact (at 20kDa, approximately three-fourths the size of GFP), tunable, and novel class of potency-enhancing protein tags for transcriptional engineering. By coupling a single targeting domain to oligomerize at specific loci, sHSP fusions offer a versatile addition to the synthetic biology toolkit for probing and programming transcriptional regulation in mammalian cells.

## Methods

### Plasmid construction

Constructs were generated by subcloning synthetic codon-optimized gene blocks (Twist Bioscience) of sHSPs, or PCR-amplified fragment from human cDNA for NGN2, into the PiggyBac vector -XL_One-GFP backbone (Addgene Plasmid #96930). This vector includes a doxycycline-inducible TRE3G promoter, and constitutive rtTA and Blasticidin resistance driven by an EF1a promoter. We modified this vector by adding a P2A-GFP or P2A-RFP downstream of the constitutive EF1a promoter. The backbone was linearized with KpnI-HF (NEB, R3142S) and SpeI-HF (NEB, R3133S), separated on a 1% agarose gel, and gel purified using Zymoclean Gel DNA recovery kit (NEB, D4007). Inserts were assembled into the backbone using In-Fusion (Takara, 639649) cloning reagents according to manufacturer’s protocols. The cloning product was then transformed into 10-beta competent E. coli (NEB, C3019H) and plated onto Ampicillin (100 μg/mL) plates and incubated overnight @37C. Individual colonies were selected and outgrown in 5mL LB + Ampicillin overnight @37C, followed by miniprep using ZymoPURE plasmid miniprep kit (Zymo, D4210) and sequence confirmation with Plasmidsaurus whole-plasmid sequencing.

### Cell Culture

HEK293T cells were cultured at 37°C, 5% CO_2_ in high-glucose DMEM containing sodium pyruvate and L-glutamine (Life Technologies, 11995) supplemented with 10% FBS (Life Technologies, A5670701) and 1% P/S (Life Technologies, 15140). To establish cell lines with transgene integration, we used the PiggyBac transposase system. Plasmids encoding PiggyBac vectors and transposase (4:1 ratio) were delivered via transfection with Lipofectamine 3000 (Life Technologies, L3000008) performed according to manufacturer’s instructions. Media was changed 18hrs after transfection, followed by 3 days of blasticidin S (Fisher Scientific, BP264725) selection (5ug/mL working concentration). Doxycycline (Sigma, D9891) was added to the media at 10 µg/mL working concentration and cells were collected 24hrs later for CUT&RUN or RNAseq.

Primary human adult dermal fibroblasts (ATCC, PCS-201-012) were cultured at 37°C, 5% CO_2_ in high glucose DMEM containing GlutaMAX (Life Technologies, 10566) supplemented with 10% FBS (Life Technologies, A5670701), 1% NEAA (Life Technologies, 11140), 1% sodium-pyruvate (Life Technologies, 11360070) and 1% P/S (Life Technologies, 15140), on TC-grade culture vessels coated with 0.1% gelatin (Sigma, ES-006-B). Cells were harvested for transfection using 0.05% Trypsin-EDTA (Life Technologies, 25300). Plasmids encoding PiggyBac vectors and transposase (4:1 ratio) were delivered using the Neon NxT Electroporation System (Life Technologies) at the following conditions: 5×10^6^ cells/ml density, T buffer, 1650V, 10 ms pulse time, 3 pulses. Transfected cells selected were with blasticidin S (5 µg/ml working concentration). Once enrichment of transfected cells was >90%, transgene expression was induced with doxycycline (2 µg/ml working concentration) and cells were collected for RNA samples after 96hrs. All downstream processes included collection of equal numbers of cells in all compared treatment groups.

### CUT&RUN

500k cells were collected and immediately processed using the Active Motif ChIC/CUT&RUN with Active Motif CUT&RUN Spike-In Control. Library preparation was performed with the NEBNext Ultra II DNA library prep kit according to manufacturer’s instructions. Library size and concentration were assessed using the Agilent Tapestation 4200 and Qubit 4 Fluorometer, respectively. Libraries were sequenced on Illumina NovaSeq X Plus (2×150bp). Reads were trimmed and filtered with fastp (0.23.4), aligned to hg38 genome with bowtie2 (v2.5.4), and peaks were called per-sample against an IgG negative control using MACS2 (2.2.9.1) callpeaks with the --broad to better capture oligomerization domains.

Overlapping peaks were extracted via bedtools (v2.30.0) intersect, and merged peaks were generated with cat. IgG normalized bigwig files were generated using deeptools (v3.5.5) bamcompare and heatmaps were produced via deeptools computeMatrix and plotHeatmap. Spike-in scaling factors were estimated in R (v4.3.2) with the DESeq2 (v1.42.0) command estimateSizeFactors using reads assigned to the dm6 genome. Spike-in normalized bigwig files were generated using deeptools bamcoverage and used for genome browser visualization on IGV (v2.16). Per-peak signal was extracted from spike-in normalized bigwig files centered on the peak using pyBigWig (v0.3.24). Binding signal correlation was assessed using Pearson correlation on log2-transformed group mean signal at +/-250bp around the peak centers. Peak center enrichment was quantified for each sample using the same +/-250bp bin and compared using a two-sample t-test. Binding breadth for each sample was measured as the average full width at half maximum (FWHM) and compared between groups using a two-sample t-test.

### RNAseq

For each replicate, 500k cells were collected and snap frozen for RNAseq. Total RNA was isolated using Quick-DNA/RNA Miniprep Plus Kit (D7003) according to manufacturer’s instructions and assessed for integrity via Agilent Tapestation 4200. Library prep of purified RNA was performed using illumina truseq stranded mrna library prep kit according to manufacturer’s instructions. Library size and concentration were assessed using the Agilent Tapestation 4200 and Qubit 4 Fluorometer, respectively. Libraries were sequenced on the Illumina NovaSeq X Plus (2×150bp). Reads were trimmed and filtered with fastp (0.23.4), aligned to hg38 genome with STAR (v2.7.9a), and counts called with featureCounts (v2.0.6).

Analysis was performed on R (v4.3.2) using DESeq2 (v1.42.0) (Benjamini-Hochberg False Discovery Rate of < 0.05). Heatmaps were generated from DEGs using scaled DESeq2 normalized data and the ComplexHeatmap package (v2.18.0). GSEA was performed using enrichplot (v1.22.0) and clusterprofiler (v4.10.0). Venn diagrams were generated with VennDiagram (v1.7.3). Scores were assigned to sHSPs by calculating the median z-scored value for up DEGs as a proxy for overall gene expression impact, and plotted using ggplot2 (v3.5.1).

### AF3 ACD Docking screen

Candidate ACD-containing proteins were modeled using a targeted docking pipeline designed to test whether the C-terminal IXI/V-like motifs are predicted to engage the ACD dimer. For each ACD protein, the predicted/annotated ACD-containing core sequence was submitted as two identical protein chains, representing an ACD dimer, and the C-terminal peptide containing the putative IXI/V-like motif was submitted as a third chain. Ten AlphaFold3 models were generated per condition. Wild-type CTD peptides were compared with matched motif-disruption controls in which the core IXI/V-like motif was replaced by alanines while preserving peptide length.

AF3 outputs were used to quantify local CTD–ACD docking. For each model, we extracted chain-pair ipTM values, peptide-to-ACD ipTM, residue-level pLDDT, predicted aligned error, and residue-residue contact probabilities. The primary docking window was defined as the IXI/V-like motif or the specified mutant residues in the CTD peptide. A model was classified as a docking hit when the motif made ACD contact with sufficient local confidence, defined by motif-to-ACD contact probability ≥0.5, local motif-to-ACD PAE ≤15 Å, and motif pLDDT ≥50. For each condition, the docking hit rate was calculated across the ten AF3 models. A composite Docking Capacity Index (DCI) was then calculated as: DCI = 0.6 × hit rate + 0.4 × mean (contact probability, PAE confidence, motif pLDDT/100, peptide-to-ACD ipTM), where PAE confidence = max (0, 1 − local PAE/20). Motif specificity was calculated as WT DCI minus the highest DCI observed among matched motif-disrupted or scrambled controls. Proteins with high WT DCI and large WT – control separation were classified as motif-dependent dockers, whereas proteins with high WT docking but persistent mutant/control docking were classified as nonspecific or noncanonical CTD–ACD contacts.

### RT-qPCR

500k cells were counted and collected for RNA extraction for each sample. Total RNA was isolated from pooled HEK cells using NEB Monarch Total RNA miniprep kit according to the manufacturer’s instructions. cDNA was synthesized from purified total RNA using the Thermo Fisher Scientific SuperScript IV VILO Master Mix (Invitrogen, 11756500). Quantitative PCR was performed using iQ SYBR Green Supermix (Bio-rad, 1708882) and gene-specific primers on a Bio-Rad CFX96 real-time PCR system. Transcript abundance was measured from biological triplicates unless otherwise indicated and analyzed using the ΔΔCt method relative to GAPDH housekeeping gene. Primer sequences used in this study are listed in Supplementary Table 2.

### Arabidopsis plant materials and growth conditions

All plants used in this study were in the Columbia-0 ecotype (Col-0, wild-type) and were grown on soil in a greenhouse under long-day conditions (16 h light / 8h dark). The mbd5/6 double mutant used in this study is the previously described mbd5/6 CRISPR (see: REF 7).

### Generation of transgenic lines

MBD6–ACD transgenic plant lines were generated by cloning codon optimized ACD sequences into an MBD6 genomic construct containing the endogenous MBD6 promoter and introns using In-Fusion cloning reagents (Takara, 639650). The assembled products were gel-purified and inserted into XbaI-digested RFP or YFP destination vectors.

Plasmids were introduced into Agrobacterium tumefaciens strain AGL0 by electroporation and transformed into Arabidopsis by Agrobacterium-mediated floral dip. Successful FLAG-tagged transformants were selected on soil using Basta resistance, whereas RFP-and YFP-tagged transformants were selected on half-strength Murashige and Skoog medium containing hygromycin

### Confocal Microscopy

Confocal imaging was carried out on a Zeiss LSM 980 confocal microscope. Arabidopsis root tips were imaged with a 40× water-immersion objective. For live-plant imaging, approximately 2-week-old seedlings grown on half-strength MS medium at room temperature, approximately 25 °C, were transferred to 1-mm-thick glass slides containing deionized water. Seedlings were positioned with the root tip centered in the imaging area and the aerial tissue extending toward the edge of the slide. A #1.5 coverslip was then carefully placed over the seedlings to avoid damaging or mechanically stressing the tissue. Typically, one to four seedlings were mounted per slide for imaging.

## Supplement

**Supplementary Figure 1).**
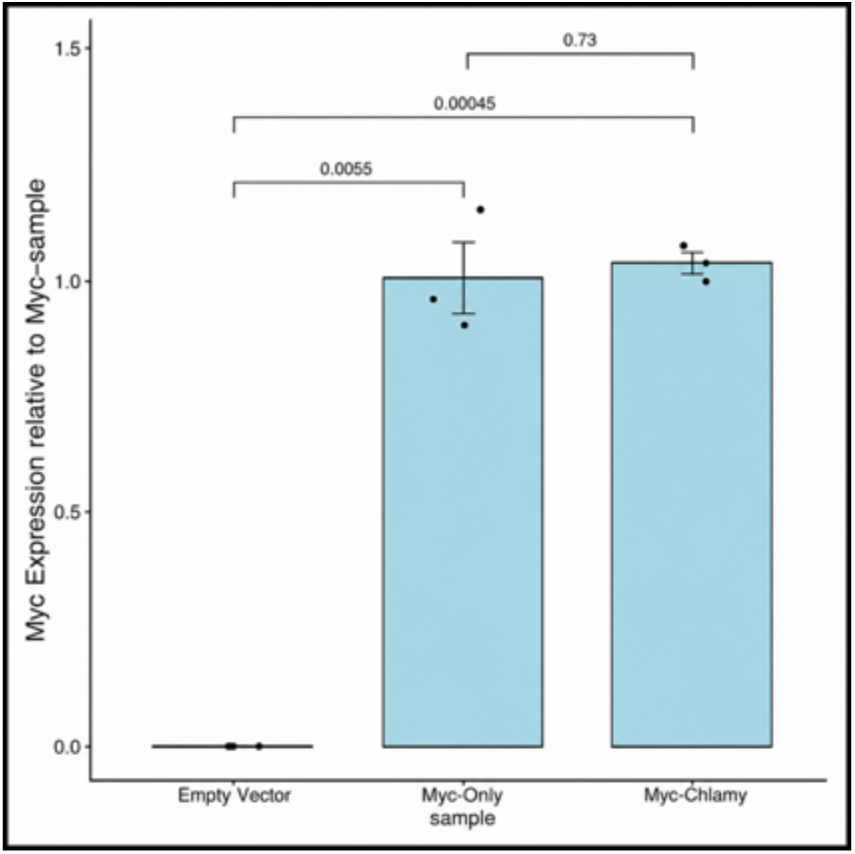
Comparable transgene RNA expression of Myc and Myc-Chlamy. RT-qPCR comparing the expression of negative control (listed as XLONE), Myc, and Myc-Chlamy HEK cell transgene overexpressing constructs at 24hrs post induction. A single pair of primers was used for both constructs, spanning transgenic Myc and the FLAG tag contained shared by both constructs.

**Supplementary Figure 2).**
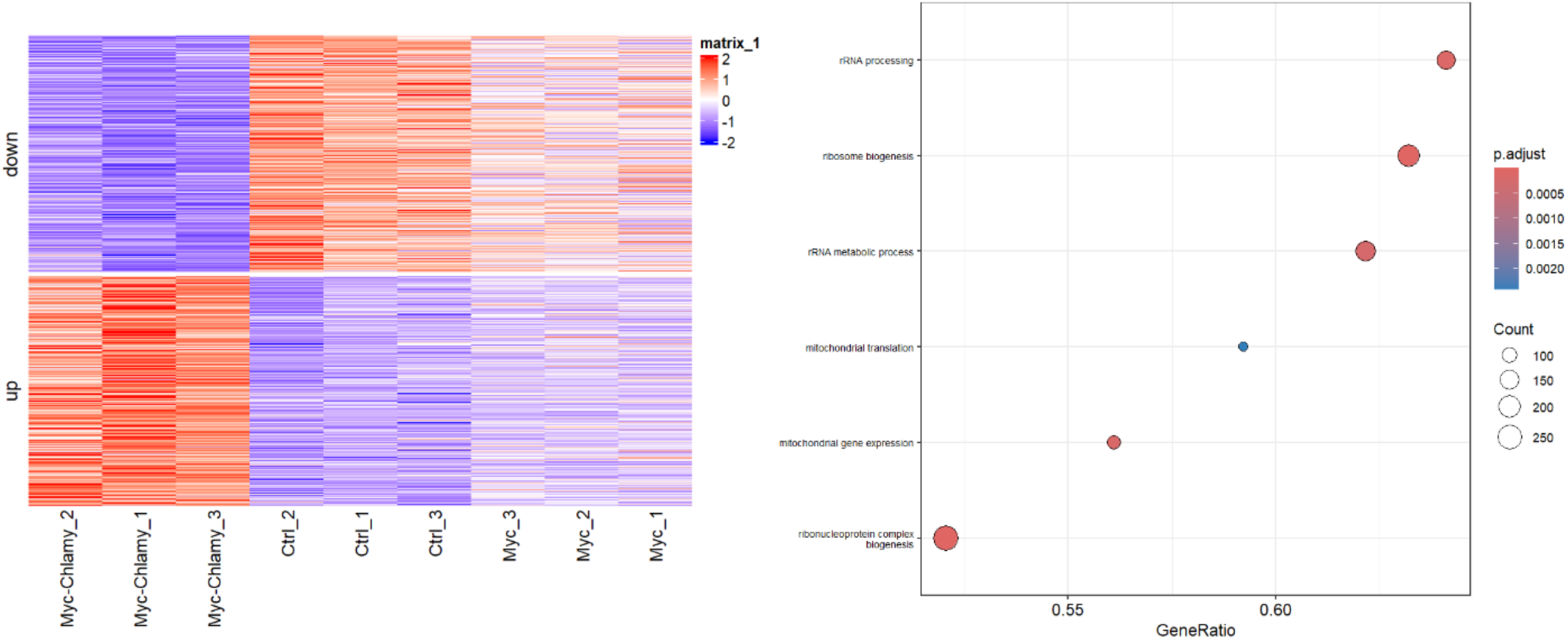
Myc-Chlamy amplifies transcriptional response associated with MYC activity. **Left)** RNAseq heatmap of differentially expressed genes (DESeq2, p.adj < 0.05) in Myc-Chlamy, but not Myc, overexpressing cells compared to control. Rows represent individual genes, columns represent biological replicates. **Right)** GSEA on genes from Left panel, highlighting biological processes associated with MYC overexpression.

**Supplementary Figure 3).**
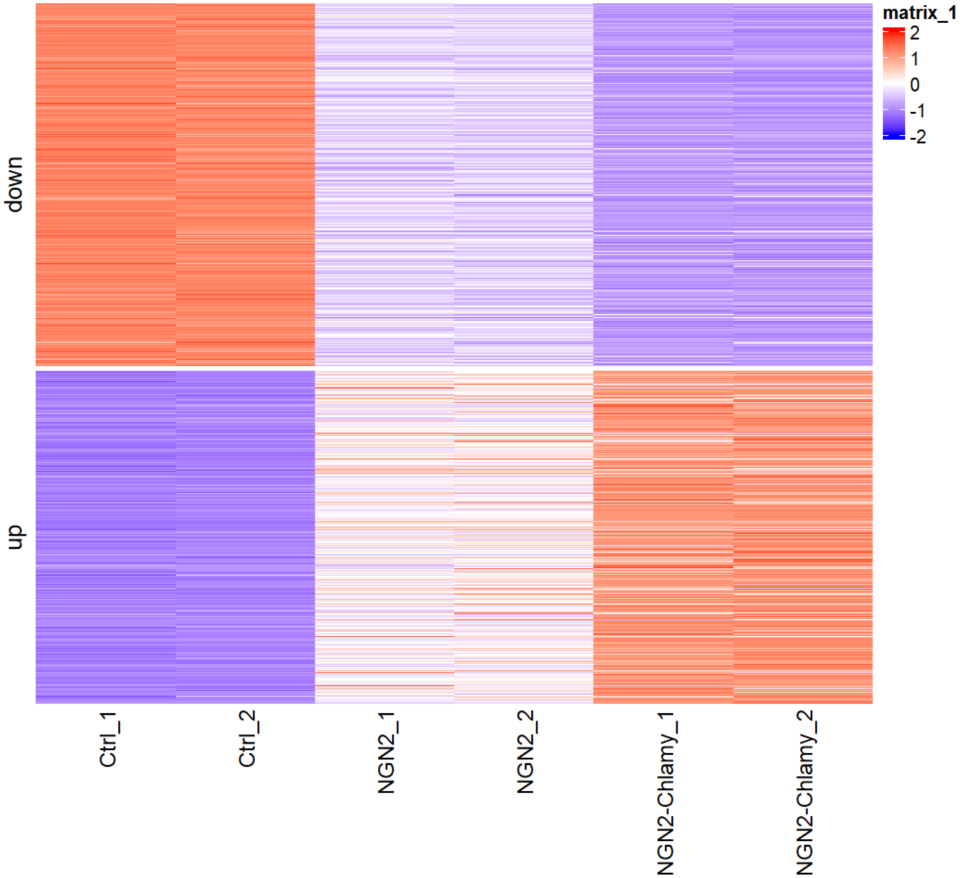
RNA heatmap showing sHSP fusion enhances NGN2-dependent transcriptional activation in primary human dermal fibroblasts. RNAseq heatmap of differentially expressed genes (DESeq2, p.adj < 0.05) in NGN2 and NGN2-Chlamy overexpressing cells compared to control at a 96-hour timepoint.

## Acknowledgements

We thank M. Akhavan, and the UCLA Broad Stem Cell Research Center BioSequencing core for high throughput sequencing. We also thank other members of the Sobreira and Jacobsen lab for assistance and helpful discussion.

## Author contributions

T.J.B., M.J.B., and S.E.J. initially conceived the project. B.B. and S.W. provided supporting conceptualization. T.J.B., M.J.B., and S.E.J. wrote the manuscript. All authors participated in manuscript editing. T.J.B., M.J.B. performed most of the experiments with assistance from S.M.L.S, Y.H. M.H., B.L., and S.S. RNA-seq experiments were carried out by T.J.B., M.J.B. with assistance from S.M.L.S. Cut&Run experiments were performed by T.J.B., M.J.B. with assistance from S.M.L.S. S.F. performed high-throughput NGS sequencing. ALL NGS sequencing processing and analysis was performed by M.J.B. Cell Culture experiments were performed by T.J.B., M.J.B., and H.L. ACDs were chosen by B.B., T.J.B., M.J.B., and M.H. Work was supervised by D.S. and S.E.J.

## Data and materials availability

The high-throughput sequencing data generated in this study have been deposited in the Gene Expression Omnibus (GEO) database under accession number PRJNA1513388. Source data are provided with this paper.

## Ethics Declaration

Ethics declaration: not applicable.

Correspondence and requests for materials should be addressed to Steven E. Jacobsen.

---

**Supplementary Table 1).**
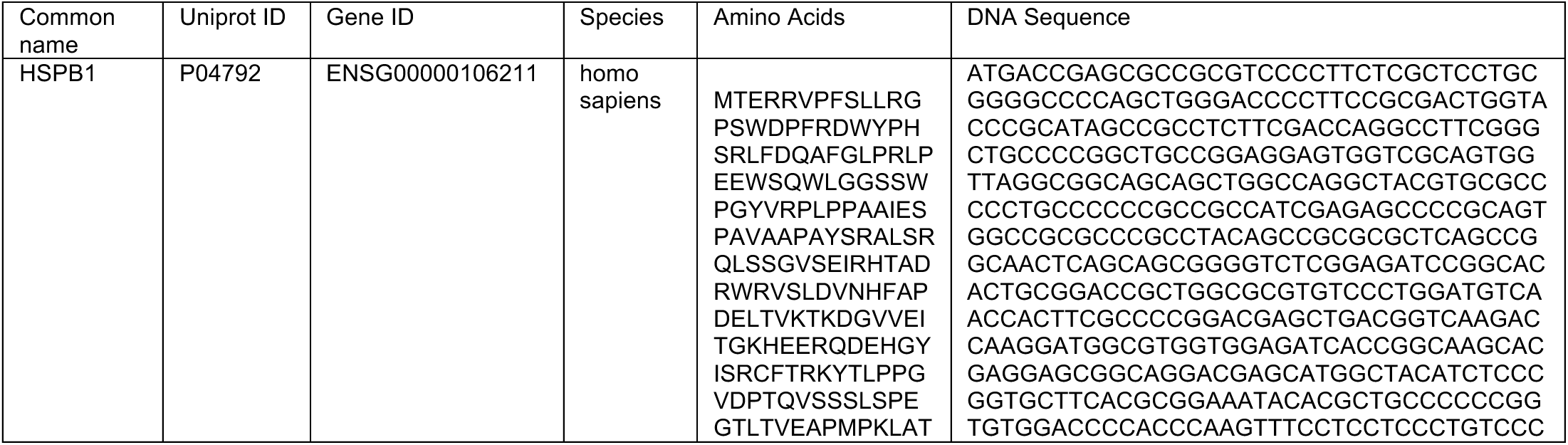
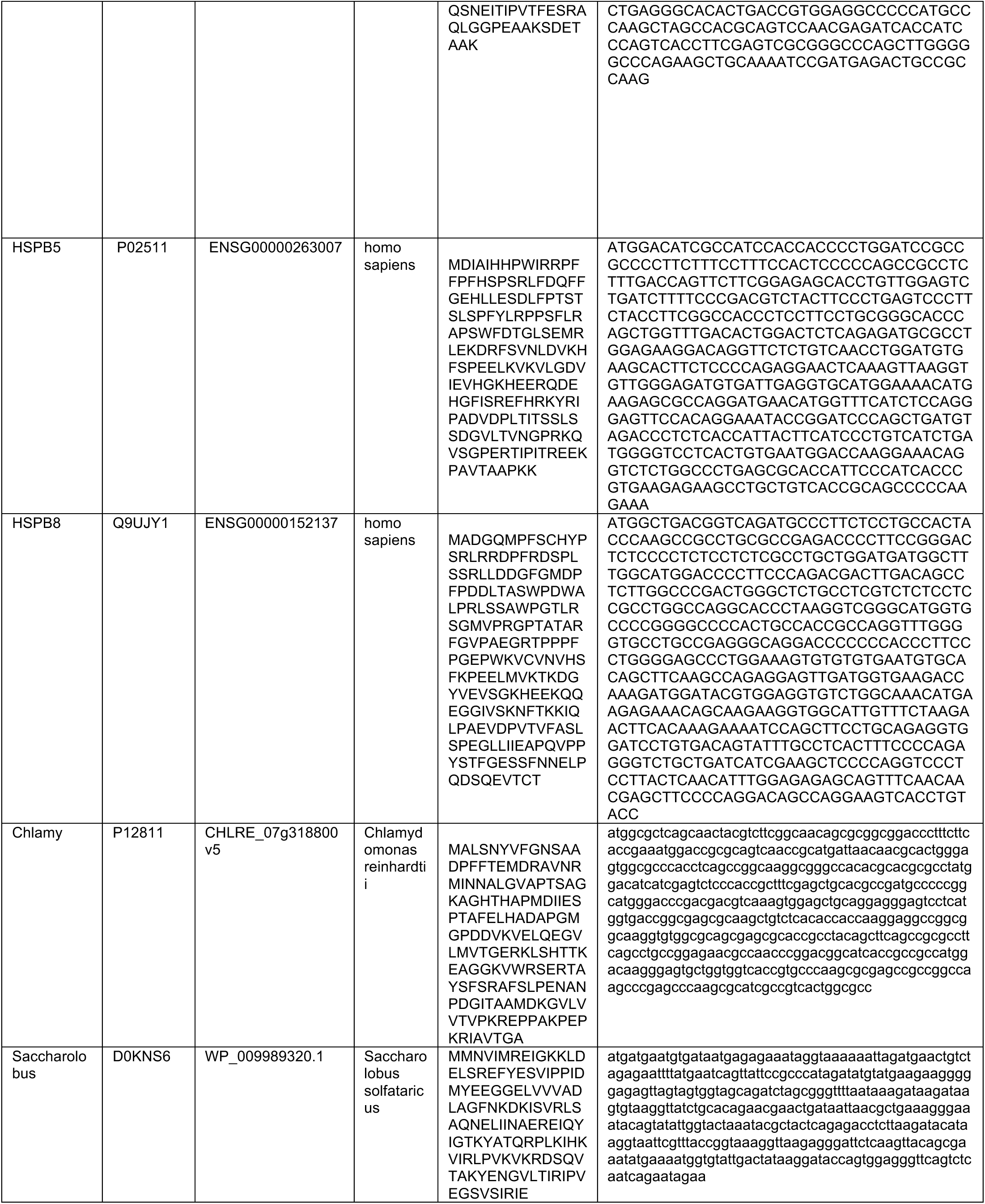
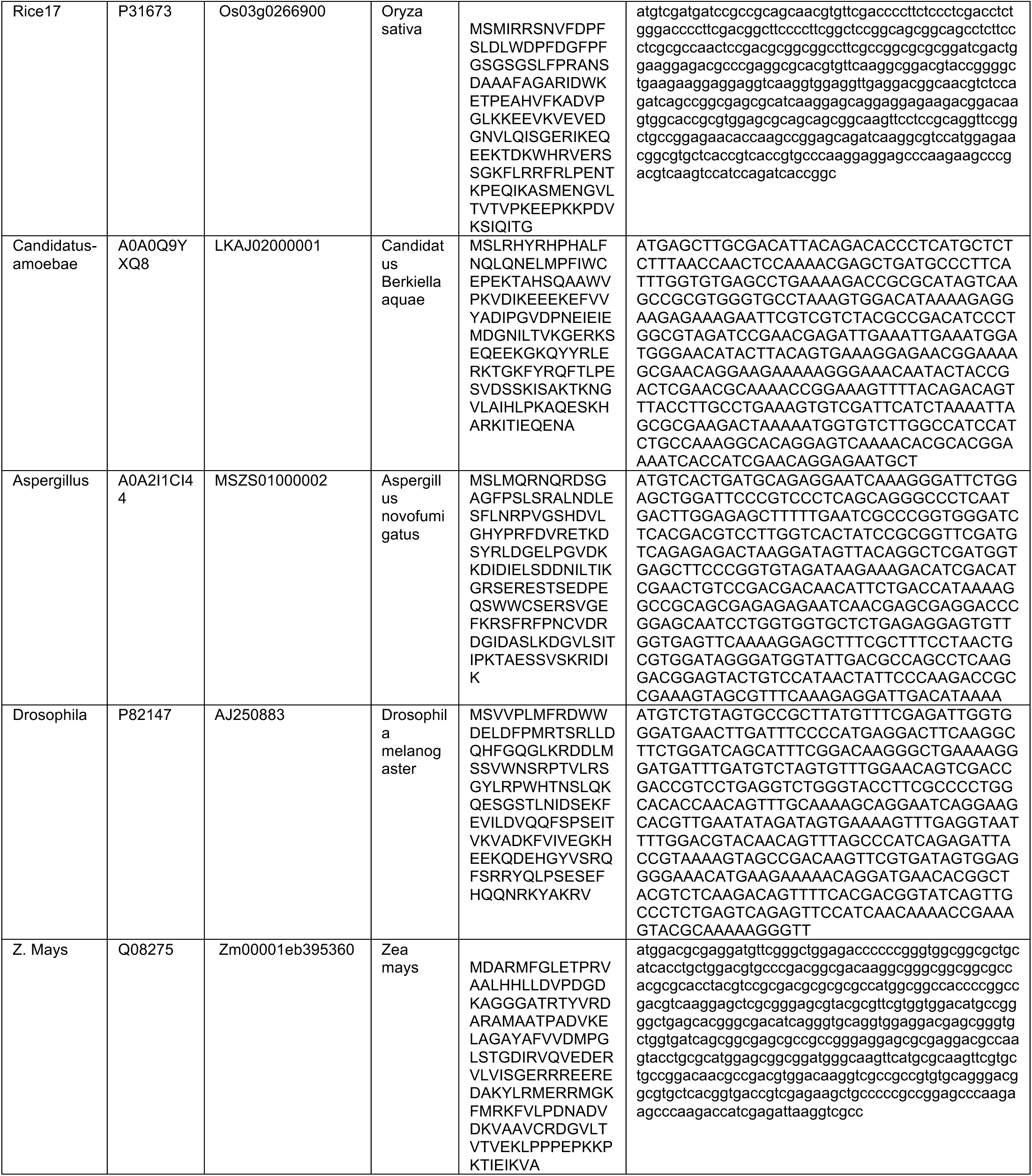
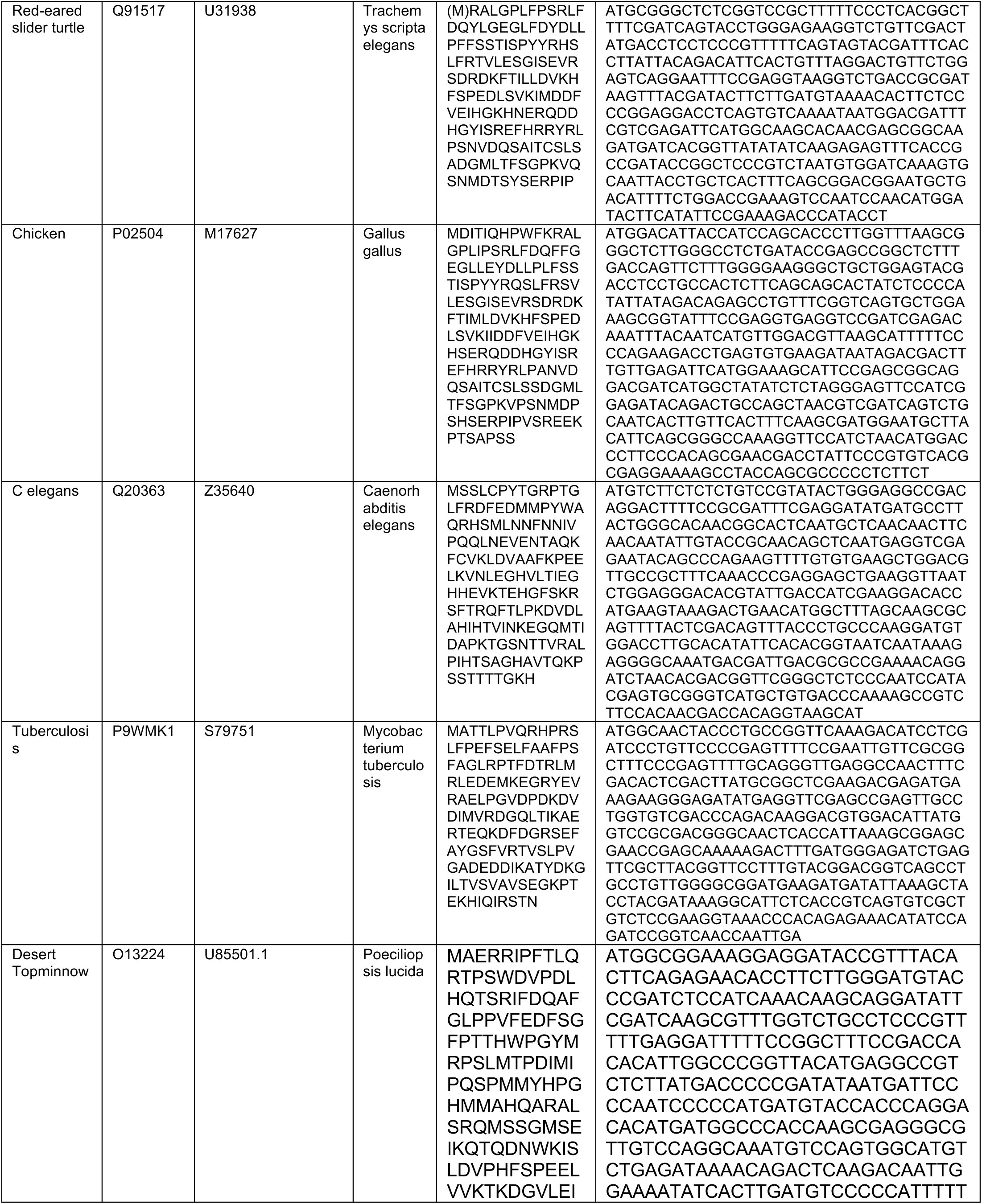
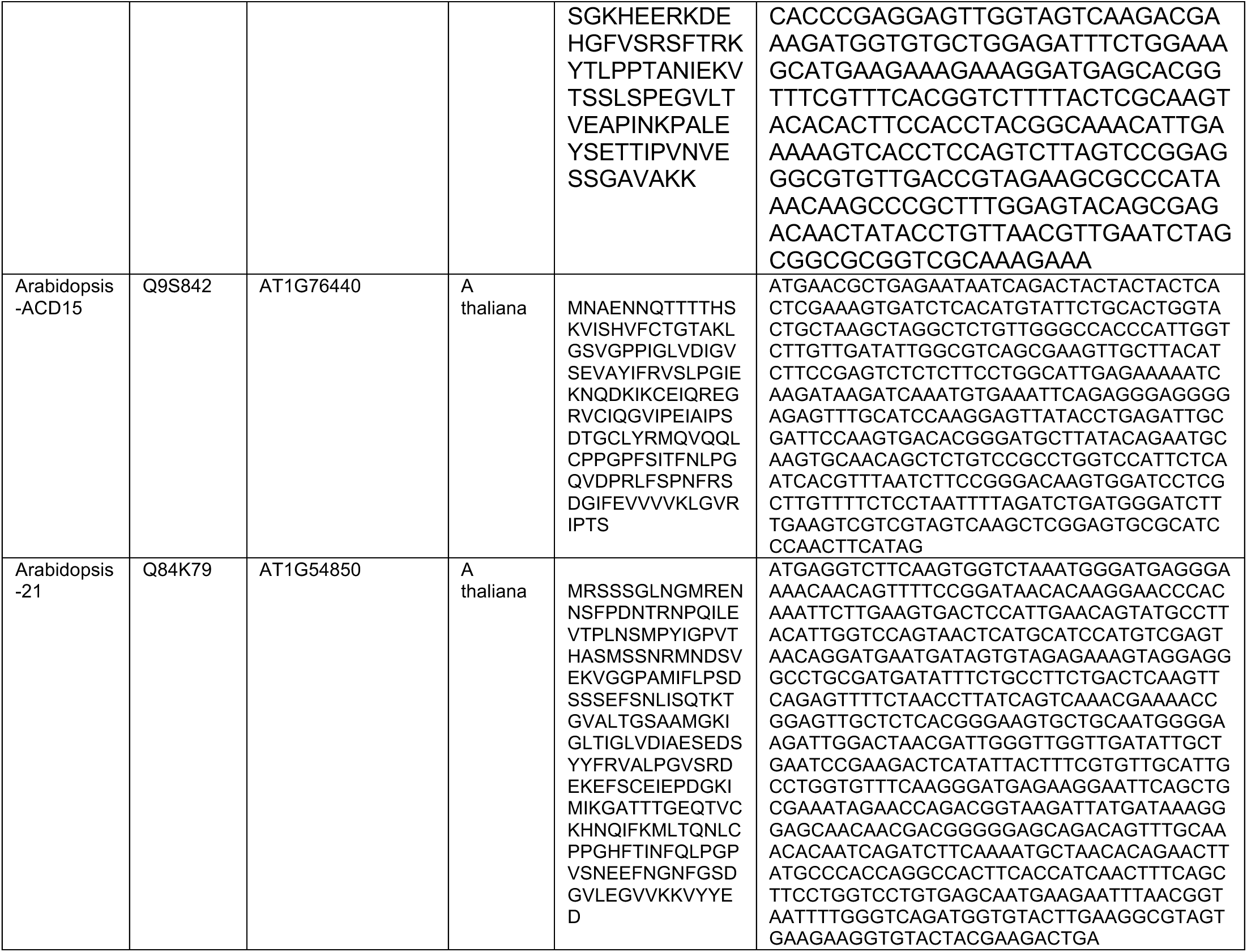
sHSPs from the tree of life.

**Supplementary Table 2).**
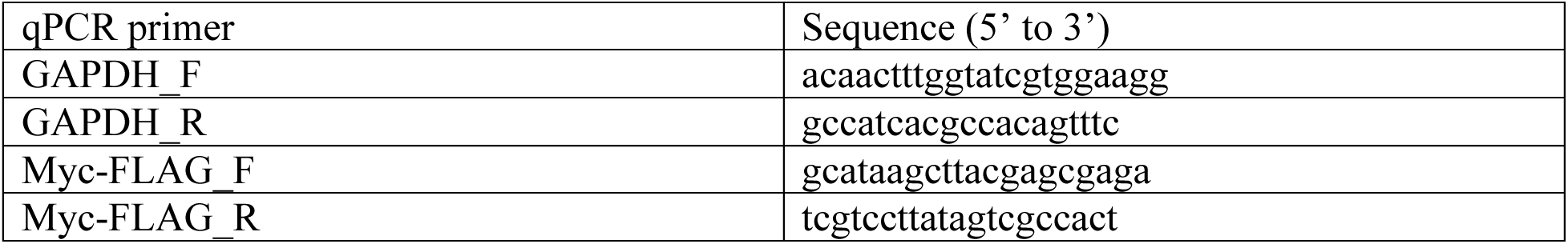
qPCR primers.

